# Neural networks simulating short-term memory of two inputs with varying commonality

**DOI:** 10.1101/2024.11.20.624539

**Authors:** Eric Raman, Larry Shupe, Ryan Eaton, Eberhard Fetz

## Abstract

The activity and connectivity of neurons in the primate brain underlying behavior cannot yet be completely specified, but neural networks provide complete models of the connectivity and activity that performs specific tasks and provide insight into the neural computations performed by the primate brain (Fetz and Shupe 2003).

Studies of neurons in the monkey cortex have shown that short-term memory of sensory events may be mediated by sustained neural activity. Short-term memory tasks have been modeled with dynamic neural networks using a single continuous variable and a gate input to create a sample-and-hold (SAH) function (Zipser 1991; Maier 2003). Networks trained to perform these short-term memory tasks develop hidden unit activity which resembles that of cortical neurons in monkeys performing memory tasks.

We here extend the investigation of single-input SAH networks to networks computing SAH for two continuous-variable inputs that have varying degrees of common mode signal. Results provide insights into computational mechanisms of associative short-term memory of sensory signals with common mode components, such as visual inputs to the two eyes, auditory inputs to the ears and proprioceptive input from multiple muscle spindle afferents.

We also examined the attractor states that these SAH networks eventually reach after sufficiently long delay periods and found that these were determined by the shapes of the input-output functions of the hidden units rather than network architecture.

## Introduction

Dynamic recurrent neural networks (Watrous and Shastri, 1986; Williams and Zipser, 1989) can be trained to perform a wide range of input-output transforms for time-varying inputs (Fetz, 1993; Fetz and Shupe 2003). Their recurrent connectivity includes feedback and cross-connections between the so-called “hidden units” located in network layers between input and output layers. The connection weights can be derived from examples of the behavior by top-down error-correction algorithms. The resulting networks provide complete solutions that perform the input-output functions insofar as all activities and connectivities are completely specified. The hidden units often exhibit activity patterns that resemble the activity of neurons in animals performing the modelled behaviors.

The recurrent connections can implement closed-loop operations that sustain responses to inputs over successive time-steps, thus providing a form of memory. Zipser (1991) first demonstrated that dynamic recurrent neural networks could simulate short-term memory tasks. The networks were trained to sample the value of a time-varying input that occurred at the time of a second gating pulse and to hold that value at the output until the next gating pulse. The activity patterns of hidden units fell into three categories: sustained activity proportional to the remembered analog value (often with a transient delay or rise), transient activity during the gate signal, and combinations of the two. These patterns were similar to the activity of neurons recorded in the cortex of monkeys performing short-term memory tasks (Fuster 1984, 1985; Funahashi 1989; Miller 1993; Rolls 2013).

We have investigated such short-term memory networks to further analyze their operation (Fetz 1993; Fetz and Shupe 2003). To elucidate the underlying computational algorithm, we constrained units to have either excitatory or inhibitory output weights (Dale 1934) with no self-connections and reduced networks towards a minimal essential network during the training process. A larger network was initially trained, then reduced by (1) combining units with similar responses and connections into one equivalent unit and (2) eliminating units with negligible activity or weak connections, then (3) retraining the smaller networks to perform the same operation. A reduced network performing the SAH function is illustrated in Figure 1. It consists of three excitatory and one inhibitory unit. The two inputs are the sample gate signal (S) and the random analog variable (A); the output (0) is the value of A at the last sample gate. This reduced network reveals a computational algorithm that exploits the nonlinear sigmoidal input-output function of the units. The first excitatory unit (SA) carries a transient signal proportional to the value of A at the time of the gate. This signal is derived by clipping the sum of the analog and gating inputs with a negative bias, as shown by the input weights to SA in the first column. This input sample is then fed to two excitatory units (M1 and M2) that maintain their activity through reciprocal connections and also feed their summed activity to the output. The inhibitory unit (SM) carries a transient signal proportional to the previous value of A. Its value is derived from a clipped sum of the gate and the previous values held in M1 and M2. As shown by its output weights, the function of SM is to subtract the previously held value from the integrating hidden units and from the output. This illustrates how the weights and activities of a complete network solution can reveal the underlying algorithm --in this case, an elegant use of non-linearity and integration to yield the appropriate remembered value. A smaller network solution exists where connections from the input units are not sign-constrained, eliminating the inhibitory unit and replacing M1 and M2 by a single self-connected M unit (Figure 1B).

**Figure 1.**
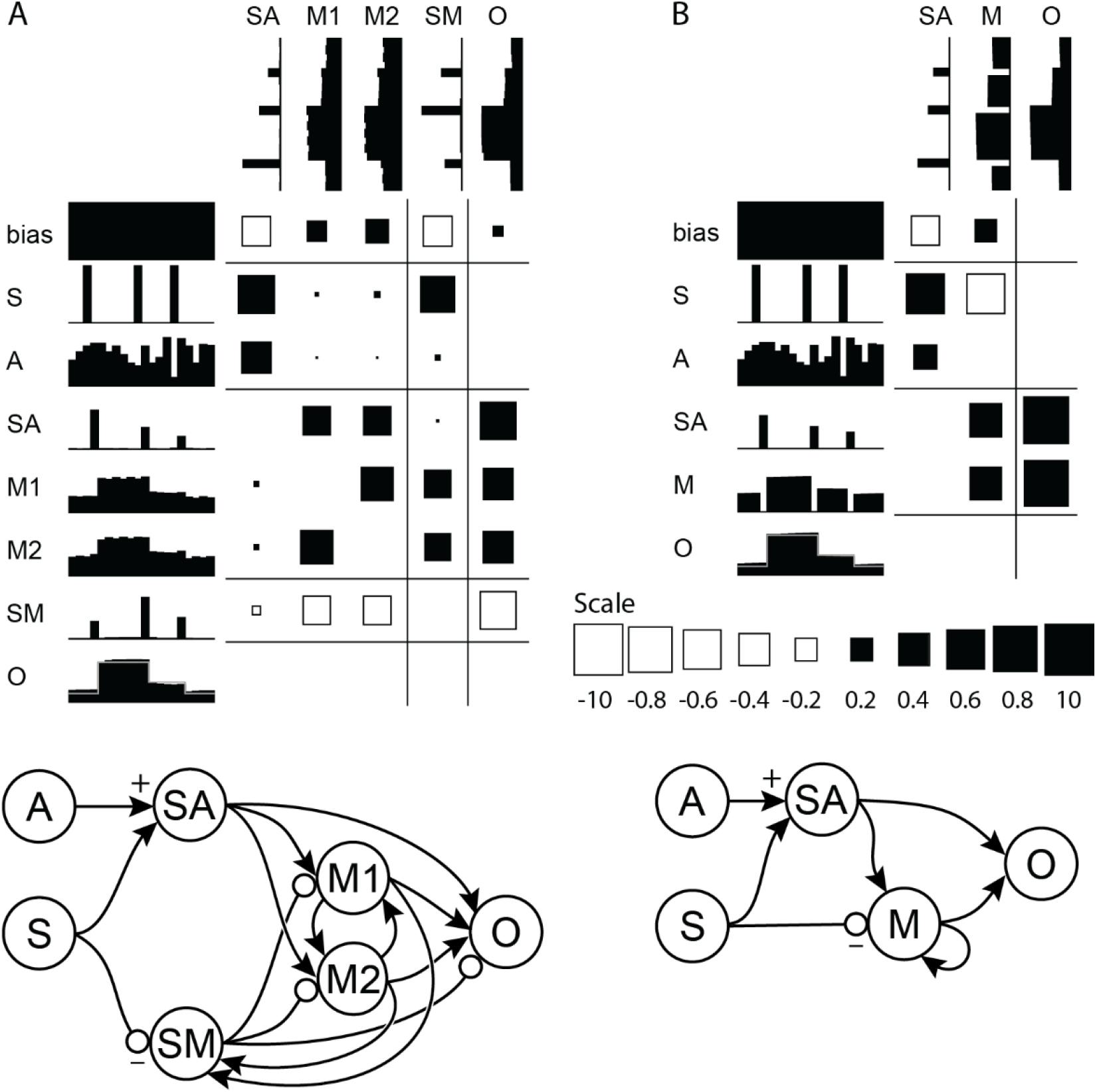
(A) Reduced network performing a sample-and-hold function, simulating short-term memory. The units are indicated by abbreviations and their adjacent activity patterns over 20 time-steps. The weights are indicated by squares (black = excitatory; white = inhibitory; scale from −10 to 10 at bottom right) with areas proportional to the connection strength from the row unit to the column unit. The two inputs are the sample signal (S) and the random analog value (A); the output (O) is the sustained value of the last sampled analog value. (B) Further reduced network demonstrating an alternative solution using a self-connected unit (M) and allowing unsigned connections from the sample input. The graph representations of the networks (bottom) show the flow of signals from input to output. Solid arrows represent excitatory connections; circles represent inhibitory connections.

The present work extends the single continuous variable input SAH networks to the case of two continuous variable input SAH networks to explore an important aspect of learning involving the recognition that multiple stimuli may have overlapping relationships with each other. In the performance of short-term memory tasks, recognition of the potential commonality between stimuli requires that the two inputs be compared. That networks can exploit the commonality is not a foregone conclusion because independent processing of the two stimuli can yield the correct output.

## Methods

We examined dynamic recurrent neural networks composed of units arranged in input, hidden, and output layers (Figure 2). Input units have activities generated by the SAH task and project unsigned (excitatory or inhibitory) connections to hidden units. Hidden units are divided into excitatory and inhibitory groups and project signed connections to other hidden units and to output units. The activities of non-input units are calculated at times *t* as *Ai(t)* = *f(Σ*_*j*_*(A*_*j*_*(t-1) W*_*ij*_*) - offset)*, where *f* is a continuous input-output activation function and Wij is the weights of the connections from unit j to unit i. The activation function is a sigmodal logistic function that saturates at a minimum value of 0 and a maximum value of 1. The offset applied to the sum of weighted activity is set to a value of 4 for these simulations and assures that units generate negligible output activity in the absence of positive input activity. Weights from input units are unsigned and constrained to the range [−8.0, 8.0], those from excitatory units are constrained to [0.001, 8.0], and those from inhibitory units to [−8.0, −0.001]. Output units are assigned target values by the SAH task. The error between these target values and the actual network outputs is computed as the square root of the mean squared difference over a series of time-steps. This error is a function of the network weights and describes an N-dimensional surface in weight-space. Neural network training is accomplished by gradient descent on this surface using a recurrent version of the back-propagation algorithm which employs both a first and second order gradient estimate as well as a line-search along those gradients to improve training speed (Watrous and Shastri 1986).

**Figure 2.**
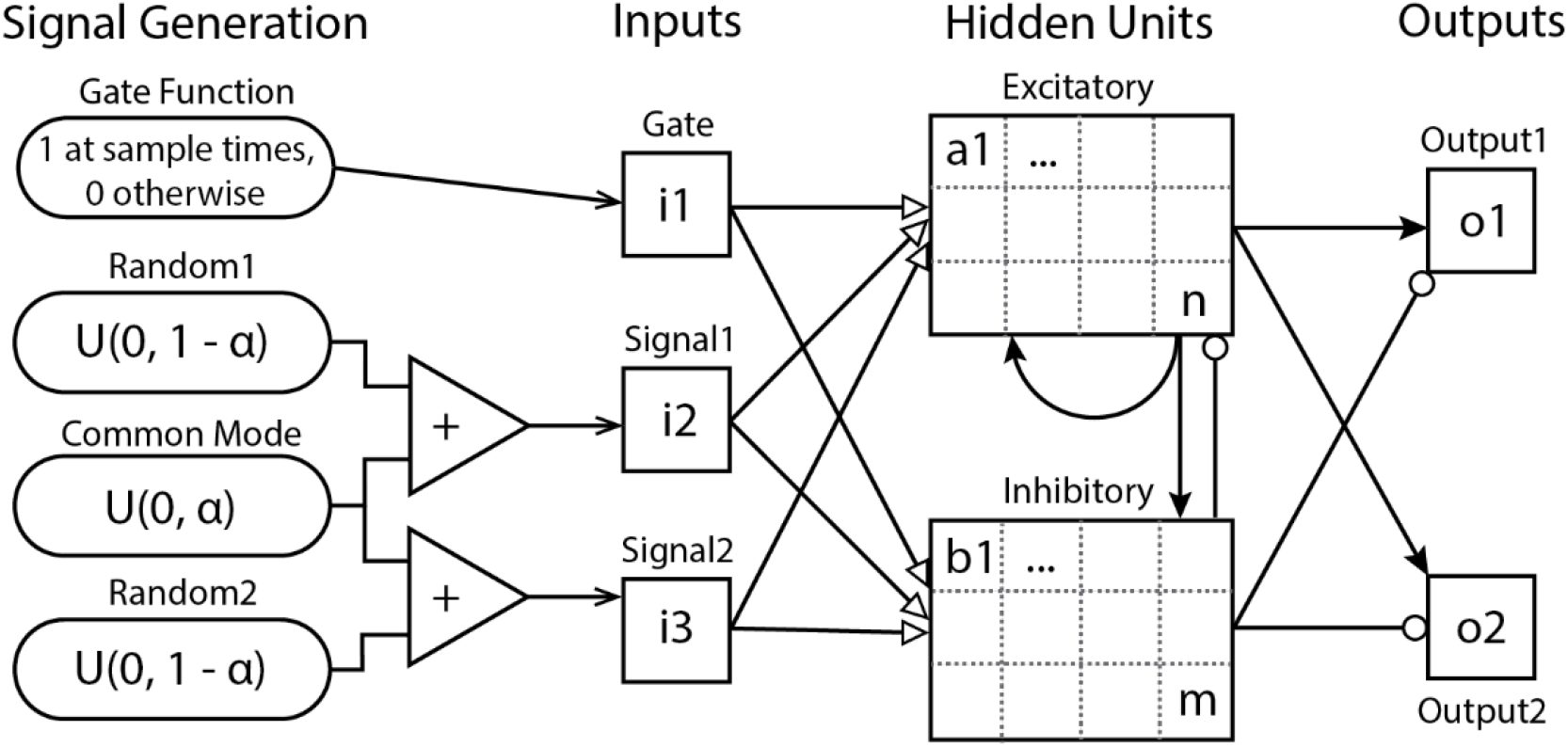
Network architecture. Input signals for each time-step are generated randomly with a Gate pulses occurring for a single time-step spaced 3 to 6 time-steps apart. Two variable Signals are created from two independent uniform random number sequences U(0, 1 – α) with an additive common mode signal U(0, α). Arrows indicate possible connections to and from all units of a category. Solid arrows represent excitatory connections and circles represent inhibitory connections. Connections from inputs are unsigned and represented by hollow arrows. Each hidden and output unit also has a single unsigned weight from a bias unit (not drawn).

Two non-independent input signals could be related to each other in several ways. One of the simplest is to extend the SAH strategy to two continuous variable inputs (Signal1 and Signal2) each sharing the same fraction (α) of a third “common mode” signal (C) as follows:

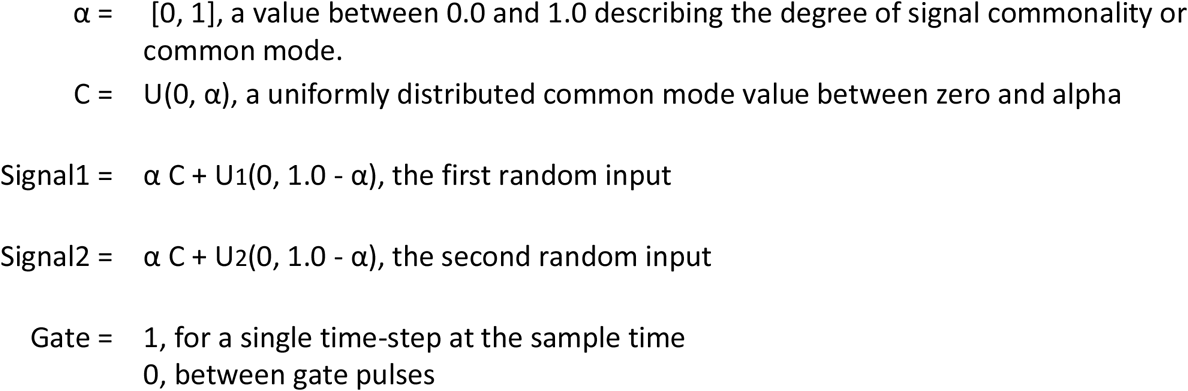

In addition to the three task-oriented inputs, a Bias input unit is maintained at a constant value of 1. Unsigned weights projecting from this unit to the hidden and output units allow the gradient decent method to adjust the activation curve offset for these connected units. The target outputs are the maintained value of Signal1 and Signal2 at the time of the previous gating signal but delayed by two time-steps to accommodate network propagation delays. It should be noted that no explicit demands are placed on the network to detect the degree of commonality between the two signals.

The held sample values and target output-values for time-step *t* are defined as:

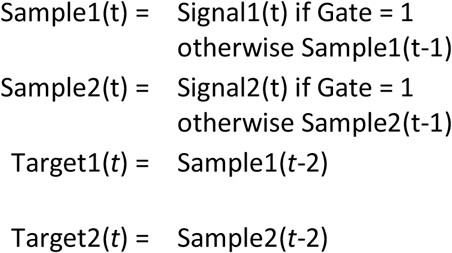

Training a large network initially helps to ensure that a solution can be found; however, to gain insight into the underlying computational mechanisms, techniques that reduce and eliminate weights and units that contribute minimally to the output are applied. Thus, training of a neural network generally involves two stages: (1) initial minimization of error in weight space, and (2) a reduction scheme used to remove minimally active and/or contributory units and to reduce unnecessarily wide distribution of activity. If the output error is kept at a low level during the reduction the underlying computational mechanism can be maintained.

Three different network reduction methods were used: weight decay, weight deletion, and unit deletion. When weight decay was active, all network weights were incrementally reduced towards zero by a specified amount (typically 0.001) after every training epoch. This allowed weights with little effect on the output to gradually decay towards zero and then be deleted when below an adjustable threshold for elimination. Similarly, units could be deleted if their maximum activity during a test period remained below a specified threshold level. After applying such network reduction, the network was retrained without weight decay to optimize the network weights for the new configuration.

All networks started with 16 cross-coupled excitatory and 16 non-cross-coupled inhibitory hidden units. Unit self-connections were not allowed and unit outputs were restricted to either excitatory or inhibitory values in accordance with Dale’s principle (Dale, 1934). Weights were initialized with random values between −1.0 and 1.0 with respect to their sign. The absence of inhibitory cross-coupled units ensures that cross-coupled memory elements will develop from excitatory units.

Networks were trained for 5000 epochs of 20 time-steps each before any weight reduction. After this initial training, three successive weight decay iterations were performed, each of which included 1000 training epochs with weight decay, followed by deletion of small weights, followed by 1000 training epochs without weight decay. These steps generated a partially reduced network for a total of 11000 training epochs. The network was then further reduced by deleting units whose maximal activity over 200 time-steps was less than (1 – α)/2 and then retraining 1000 epochs with no weight decay and then 500 epochs with weight decay at which point small weights were once again deleted. The unit deletion was then performed two additional times with 500 retraining epochs after each subsequent deletion for a total of 13500 training epochs in the fully reduced network. The network configuration was saved at specified intervals with the three most relevant networks saved at training epoch 5000 (unreduced network), epoch 11000 (partially reduced network), and epoch 13500 (reduced network). During all weight reduction iterations, a line-search along the first order gradient was used if the resulting error was less than that achieved for the second order gradient. Networks were trained for common modes between 0% and 100% in increments of 10%.

## Results

Figure 3 shows a network trained at 0% common mode, with units arranged according to their contribution to the two output signals. The full-sized network is the unreduced version after 5000 training epochs. The network size was gradually decreased from 32 hidden units and 946 weights to a final size of 8 hidden units and 49 weights (Figure 4) with minimal increase in error (from 2.93% at 5000 epochs to 3.73% at 13500 epochs [Table 1]). Comparison of all three networks shows that the underlying structure has been retained: units a5 and a7 function as memory cells for Signal1, while a6 and a14 function similarly for Signal2. Units a5, a7, and a14 are cleared by the Gate, which also saturates a6. Unit b15 generates a graded pulse proportional to 1 – Signal2 (i3) at the Gate. This then is subtracted from the second integrator and Output2. The sub-network driving Output2 is completely decoupled from Signal1, while the sub-network for Output1 receives input from both signal inputs primarily through a12, a15, and b15, although most of the activity traces back to a combination of the Signal1 and the Gate as removing all weights from Signal2 has very little effect on Output1 even though there are modest weights from a6 and a15 to Output1. Thus, the network has divided itself into two predominantly independent sub-networks.

**Table 1.** Average percent error (over 100 epochs of 20 time-steps each) and network sizes throughout the training process for the 0% common mode network.

| Training Epoch | # Hidden units | # Weights | % Error (n=100) |
| --- | --- | --- | --- |
| 0 | 32 | 946 | 54.05 |
| 3000 | 32 | 946 | 3.08 |
| 5000 | 32 | 946 | 2.93 |
| 6000 | 32 | 946 | 2.43 |
| 7000 | 32 | 648 | 2.32 |
| 8000 | 32 | 648 | 2.65 |
| 9000 | 32 | 483 | 2.61 |
| 10000 | 32 | 483 | 4.70 |
| 11000 | 11 | 70 | 3.54 |
| 12000 | 8 | 51 | 3.97 |
| 12500 | 8 | 51 | 4.28 |
| 13000 | 8 | 49 | 3.90 |
| 13500 | 8 | 49 | 3.73 |

**Figure 3.**
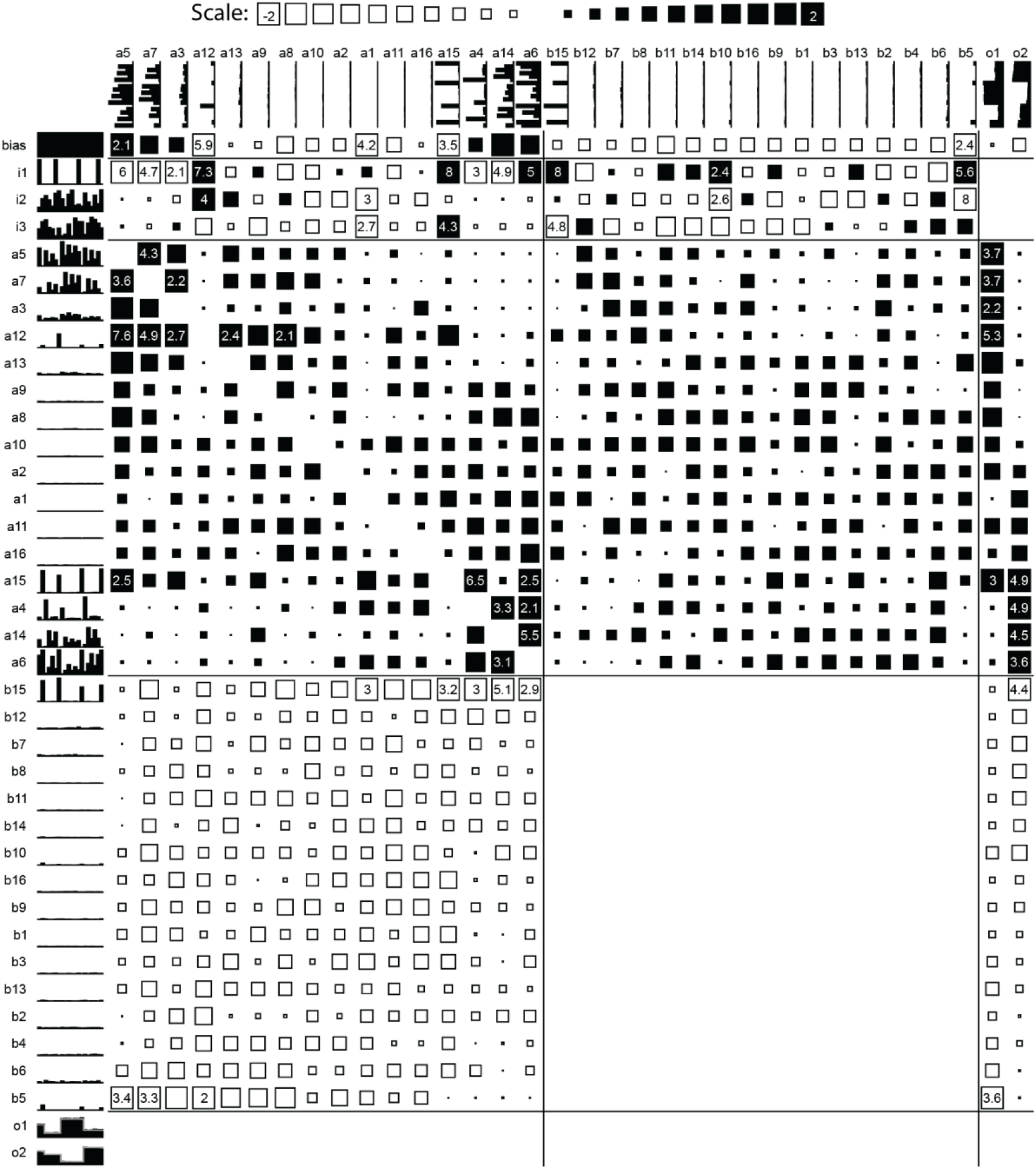
Unreduced network trained for 5000 epochs with 0% common mode inputs. Connectivity matrix shows size of excitatory (black) and inhibitory (open) weights scaled up to value of 2 and given numerically when above that. Blank spaces are unused connections. Two random inputs (i2 and i3) are sampled at the times of gating signal (i1) and those values, delayed by 2 time-steps, are presented as the target for outputs o1 and o2 (grey traces over the output values). Hidden units are ordered by their relative contributions to o1 and o2.

**Figure 4.**
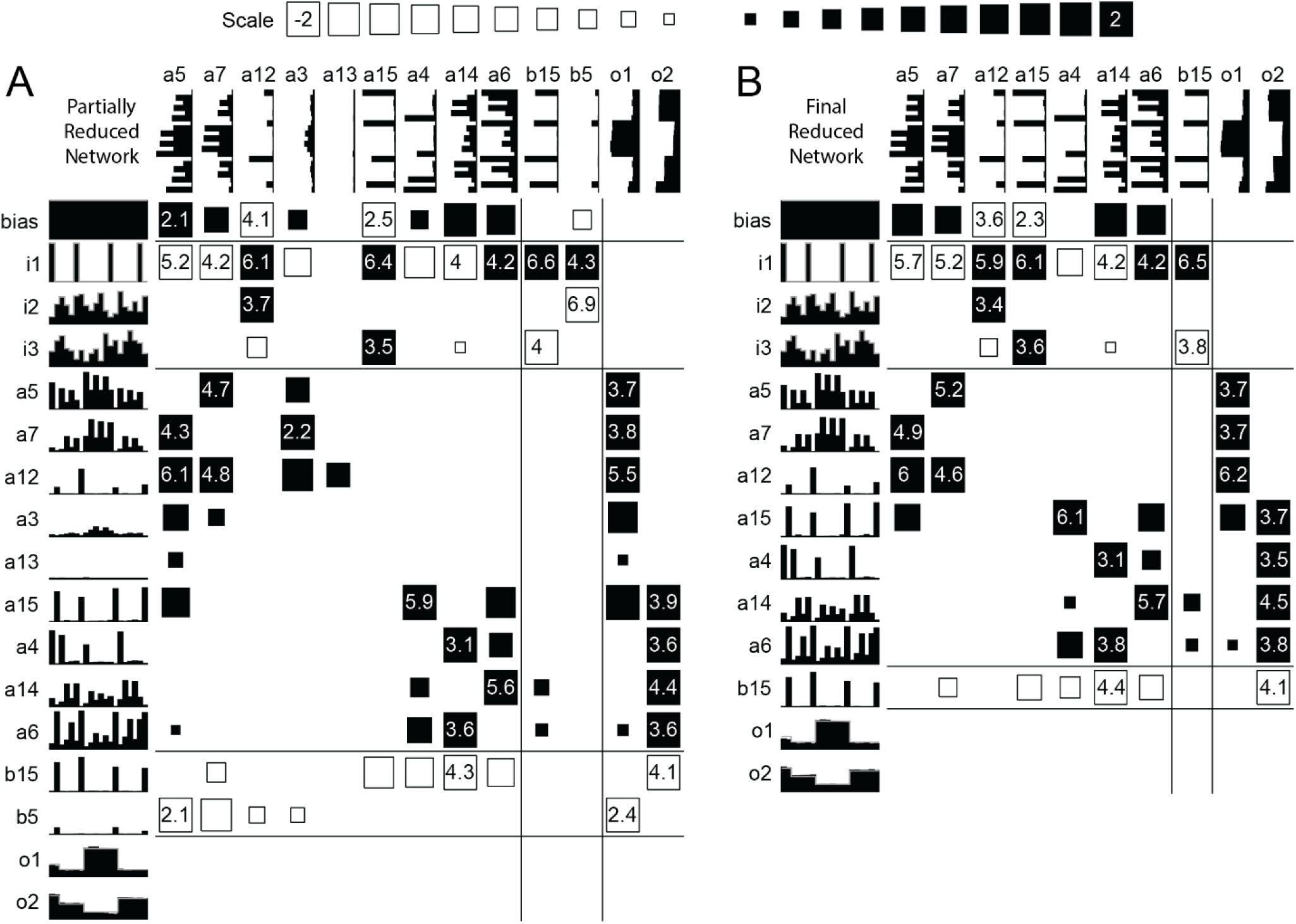
Reduced networks trained with 0% common mode inputs. (A) Partially reduced network (11000 epochs). (B) Final reduced network (13500 epochs)

The performance of networks trained with 0% common mode and tested with varying amounts of common mode is given in Table 2. As expected, the 0% common mode solution performs well across the range of common mode, because the two continuous variable input signals are essentially processed independently. The final error tends to be slightly smaller for 10% to 90% common mode values partially because these will produce fewer extreme target values (near 0.0 or 1.0) which are difficult to achieve due to the nature of the logistic activation function.

**Table 2.**
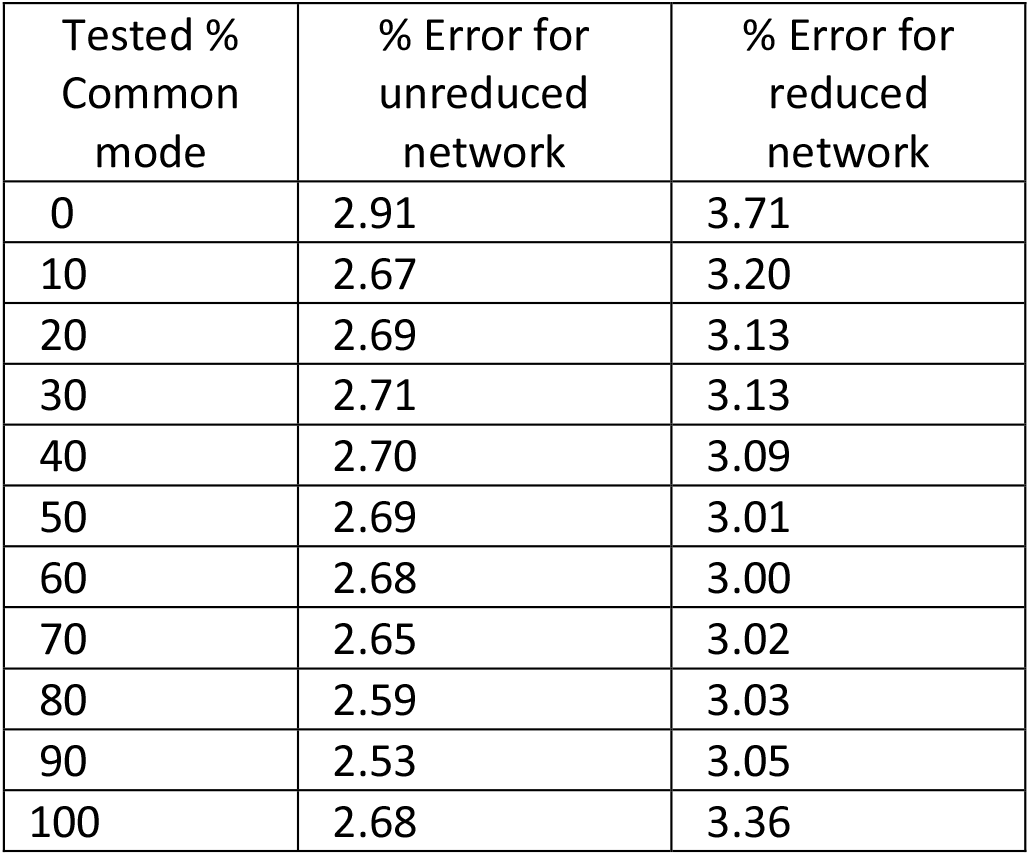
Average percent error (over 100 epochs) for the 0% common mode network when tested with other levels of common mode.

Other networks trained on common mode inputs greater than 0% showed that the basic approach of the unreduced network was retained in the final reduced network. Because the networks were reduced only via the automated reduction scheme, some further reduction is possible via careful manipulation, but this was not done here.

Networks produced by the reduction process demonstrate three different basic solutions: [1] networks processing the two signals largely independently, like the 0% common mode solution described above, [2] networks processing the two signals as common and differential mode, and [3] networks processing a single average signal.

The first type of network is seen in networks trained on 0%, 10%, 20%, and 30% common mode input signals where at least one output unit has developed a sub-network that is disconnected from that of the other output unit. In the 30% example (Figure 5) Output1 is nearly decoupled from Signal2 except for a small weight between the integrator unit a15 for Output2 and the integrator unit a3 for Output1. Output2 is nearly decoupled from Signal1 except for a connection from unit a12 which receives input for Signal1 at the time of the Gate. However, testing by lesioning the weights from just one signal reveals that these connections have little effect on the held value of the other signal.

**Figure 5.**
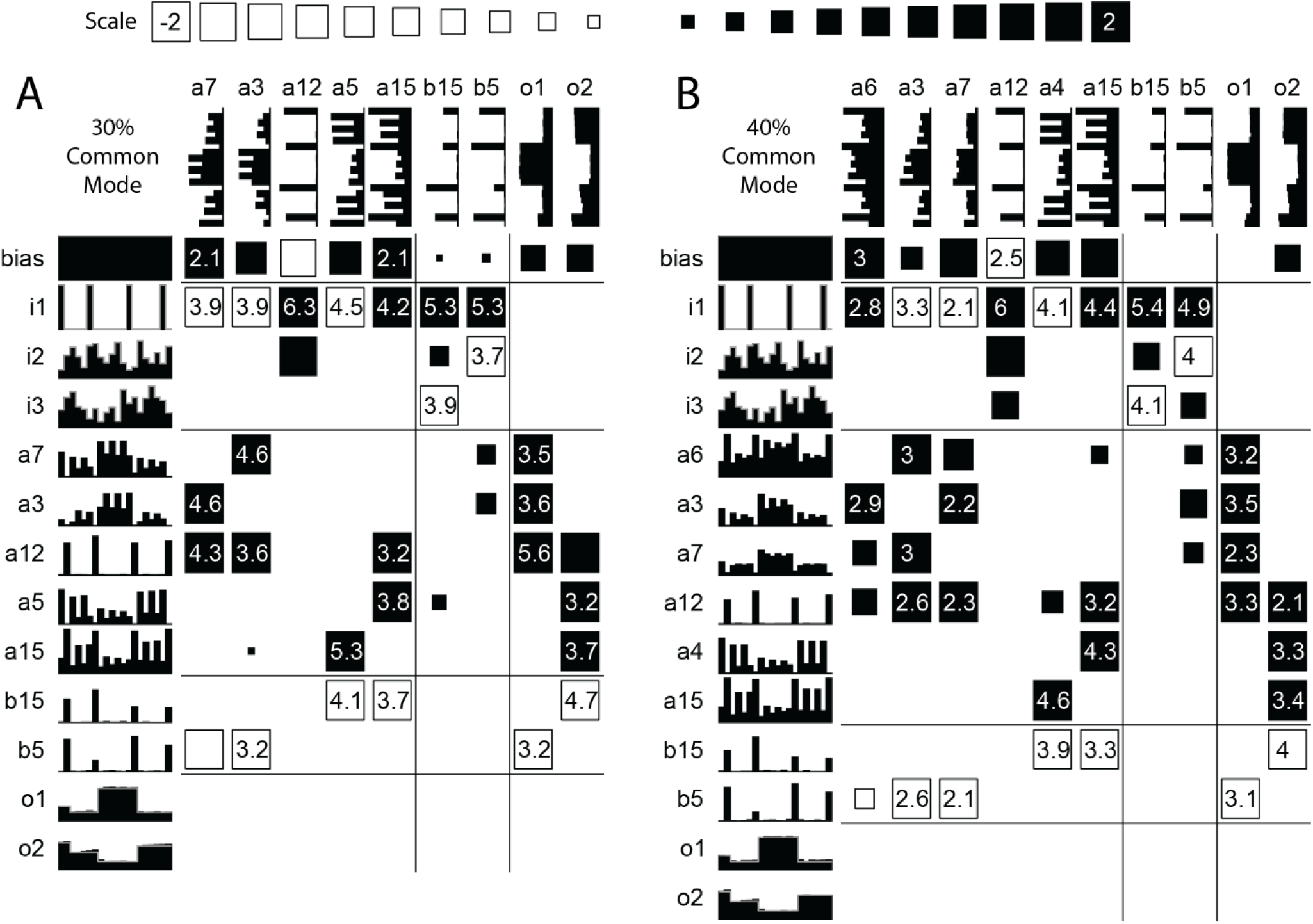
Reduced Networks. (A) At 30% Common Mode (and below) the sub-networks for Signal1 and Signal2 are nearly decoupled. (B) At 40% Common Mode, cross-coupled weights for the two input signals exist.

The second type of network appears in the 40%, 50%, and 60% common mode solutions, and to some extent in the 70% common mode solution. For example, the network in Figure 5B has one excitatory unit (a12) that derives a common-mode signal by adding the two time-varying inputs plus the gate pulse and using the negative bias to clip the peak of this sum. This transient signal is passed to both sets of cross-connected memory units (a3, a6, a7 for output1, and a4, a5 for output2) that sustain held values between gate pulses. Two inhibitory units (b5 & b15) receive differential inputs from the two input signals. Each differential input supplies the necessary modulation for the respective memory elements by removing the opposing signal’s contribution to the average estimator and emphasizing the aligned signal’s contribution. Figure 5 shows the subtle difference between the 40% network and the 30% network where outputs became more decoupled.

The third type of network does not employ the differential mode processing of the signals, and merely processes the combined signal. While the 70% common mode solution has elements of this structure, the 80%, 90%, and 100% common mode trained networks primarily process the average signal (Figure 6). There is a moderate decrease in the number of weights and hidden units remaining at the end of the reduction scheme for networks trained at 80% common mode and above compared with networks trained at 50% to 70% common mode.

**Figure 6.**
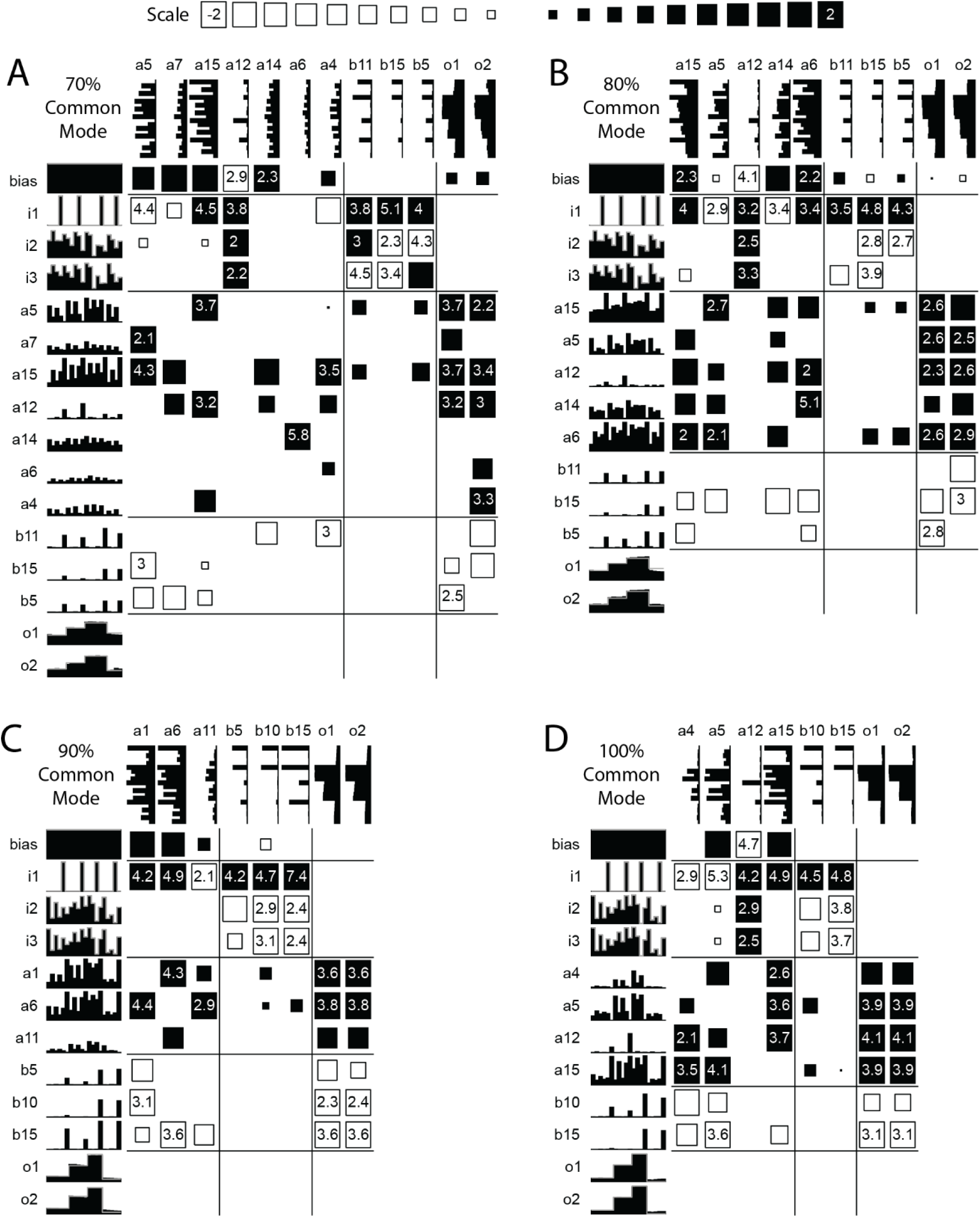
Reduced networks trained at high levels of Common Mode. (A) 70% Common Mode. (B-D) Reduced networks trained with 80% to 100% Common Mode show little differentiation between the input units. The connections to Output1 and Output2 are also very similar in these cases as the network essentially treats the input signals as the same.

These transition points at which networks change from treating the input signals as different, similar, or the same based on the level of common mode can be visualized in Figure 7, which shows how the mean output error changes over the first 5000 training iterations. These transition points are not fixed and can be changed by increasing the length of the input patterns to give the networks more information for each iteration. For the short input pattern length used here, the networks with common mode between 0 and 30% all show a smooth decent to a minimal error value (Figure 7 gray and yellow lines). Between 40% and 60% common mode, the networks have increasingly more difficulty distinguishing the two input signals, but eventually do so and continue to reach similarly low output errors (Figure 7 blue lines). For 70% common mode and above, the networks quickly settle on approximating the outputs with an intermediate value (Figure 7 green lines). The 100% common mode is an exception here since the same signal is used for both inputs and outputs. It compares closely with a network trained on a single signal input and output.

**Figure 7.**
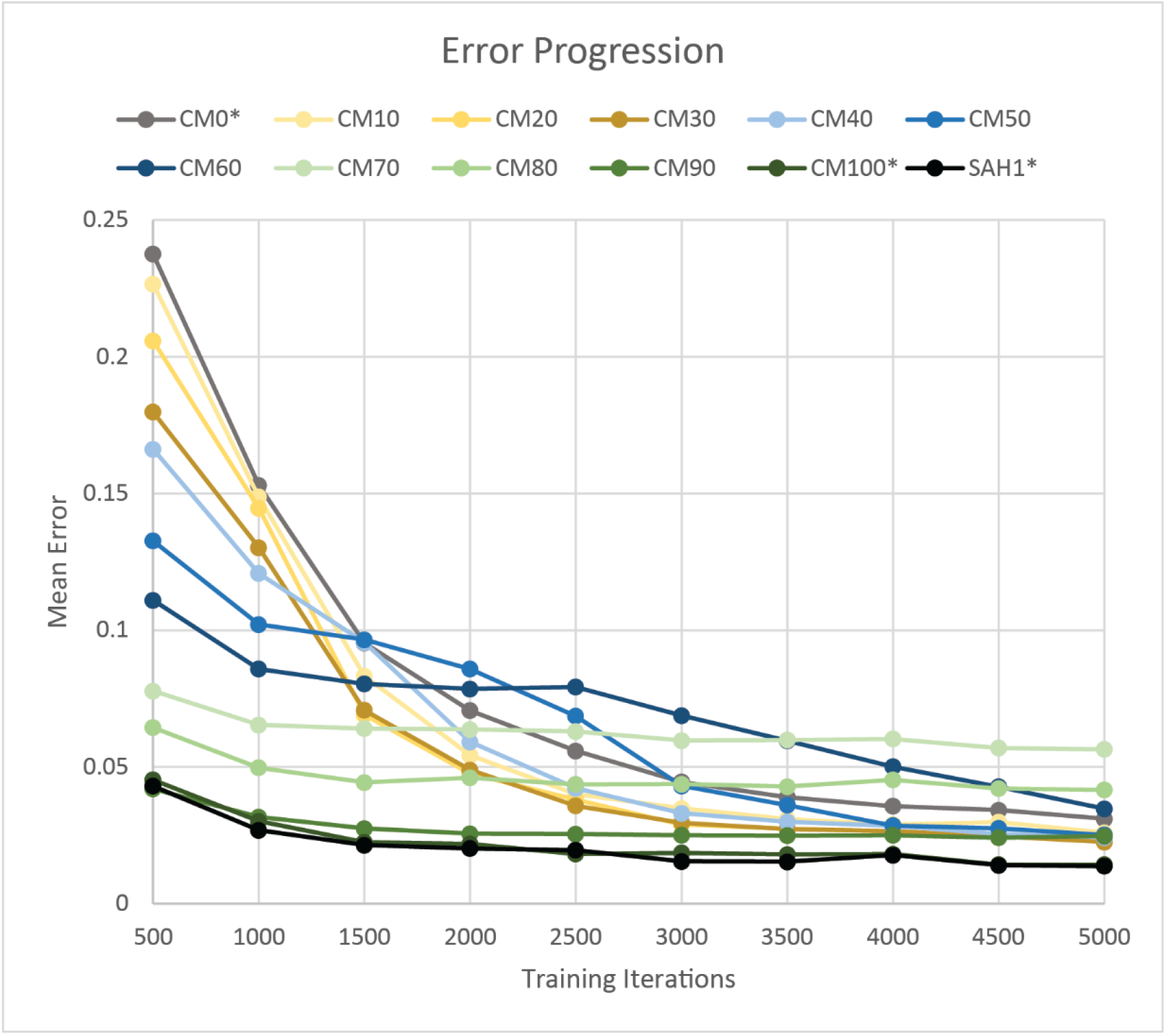
Mean error progression over the first 5000 training iterations for all tested levels of common mode (CM0 to CM100) and for a single signal SAH network (SAH1). Each point is the mean over 25 randomly initialized networks of 100-point error-averages calculated after running the appropriate number of training iterations (in steps of 500 iterations). The inputs for CM0*, CM100*, and SAH1* were slightly modified to have the same random distribution of values as CM10 and CM90 while retaining their correct level of common mode to them more comparable with the other CM levels whose signals are the sum of two random values.

Figure 8 shows the error curves produced when networks that were trained at various degrees of common mode are then presented with patterns spanning the range of common mode. Figure 8A is the error curve for the unreduced networks and Figure 8B shows errors for the reduced networks. These two curves have similar shapes, which is expected given that the reduction scheme has retained the underlying mechanisms of the unreduced networks, but also indicates that the reduction process itself is not causing disproportionate amounts of error.

**Figure 8.**
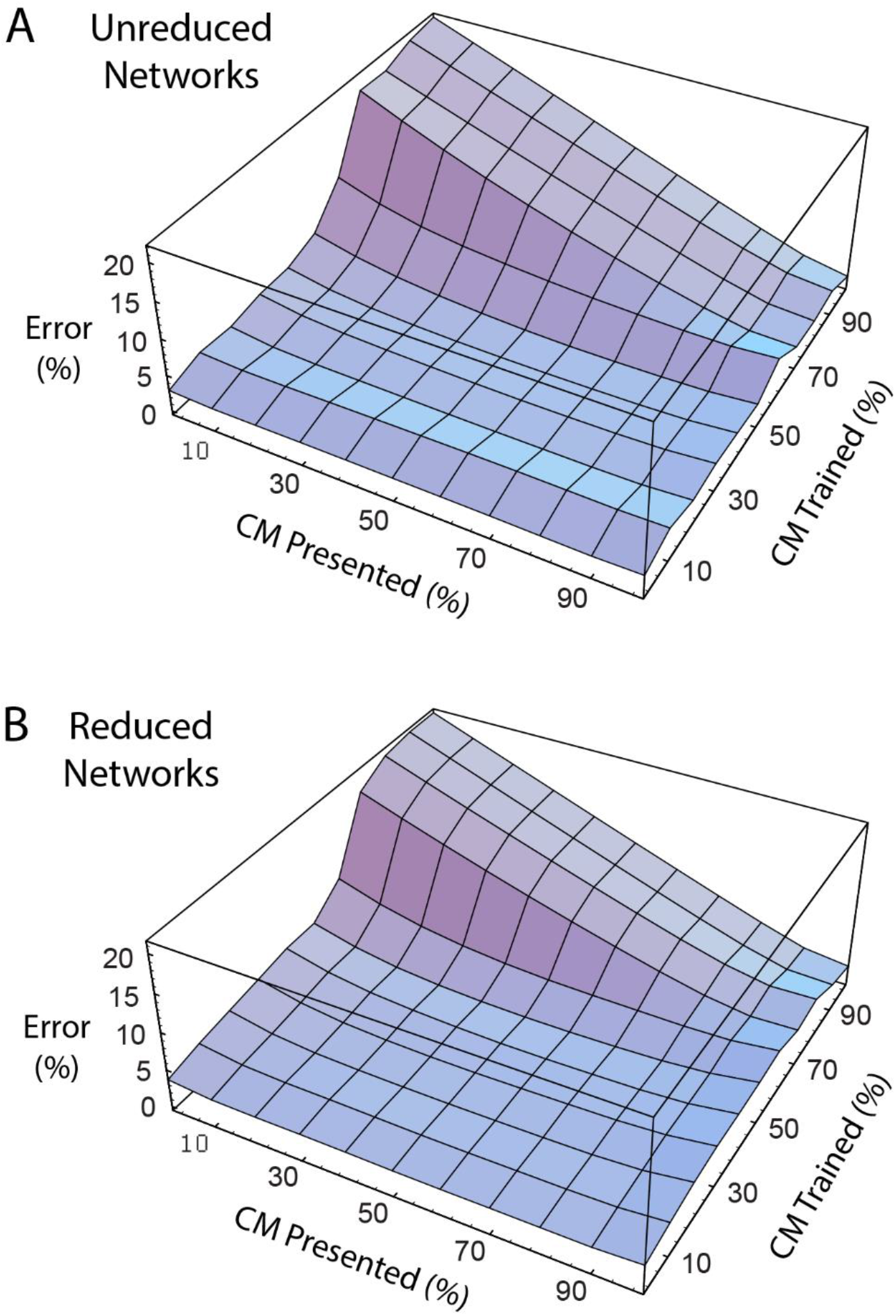
Network error for non-trained common mode input signals. A) Unreduced and B) reduced networks show similar output errors when presented with input signals on which they were not trained.

The curves show a dramatic rise in the error for networks trained at 70% or more common mode when presented with low common mode input patterns. This corresponds to the development of the third type of network described above -- i.e., networks trained at high common mode that no longer differentiate between the two inputs or the two outputs. These networks generate an average signal and split the error between the two output signals. When they are presented with low common mode inputs, the errors produced barely improve on the 29% average error expected if both outputs generated the mean of the two target outputs (or a steady 0.5 value) with 0 common mode input signals. This 29% average expected error is calculated as 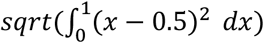.

Networks trained at or below 65% common mode produced error values of less than 5% for the degree of common mode they were trained on; the error rose slightly with presentation of patterns with common mode values either above or below that which the network was trained on. However, for networks trained at 70% common mode and above, lower error was produced when the networks were presented with patterns having greater common mode than what the network was trained on. Indeed, the highest error generated at the end of training was with the 70% common mode network at above 7% error. The same network produced only 4.2% error when presented with a 100% common mode pattern. This suggests that these networks treat both inputs as if they carried the same signal with any difference between the inputs treated as noise.

## Attractors

These SAH networks do not hold the sampled values indefinitely. Cortical firing patterns of short-term memory cells in monkey cortex (Fuster 1984, 1985) and SAH network models (Zipser 1991) both exhibit relaxation to fixed-point attractors -- specific stable activity levels to which neural networks relax during prolonged retention of information. We investigated two factors that might govern the fixed-point attractor dynamics exhibited by single-input SAH network models: network architecture and single unit input-output functions (Eaton 2007). Figure 9 illustrates the relaxation to fixed-point attractors for the reduced network shown in Figure 1B. After the end of sample pulses the values in the hidden unit and output unit decay to asymptotic values. There are two attractors (grey and black), depending on whether the last stored value is above or below a critical threshold.

**Figure 9.**
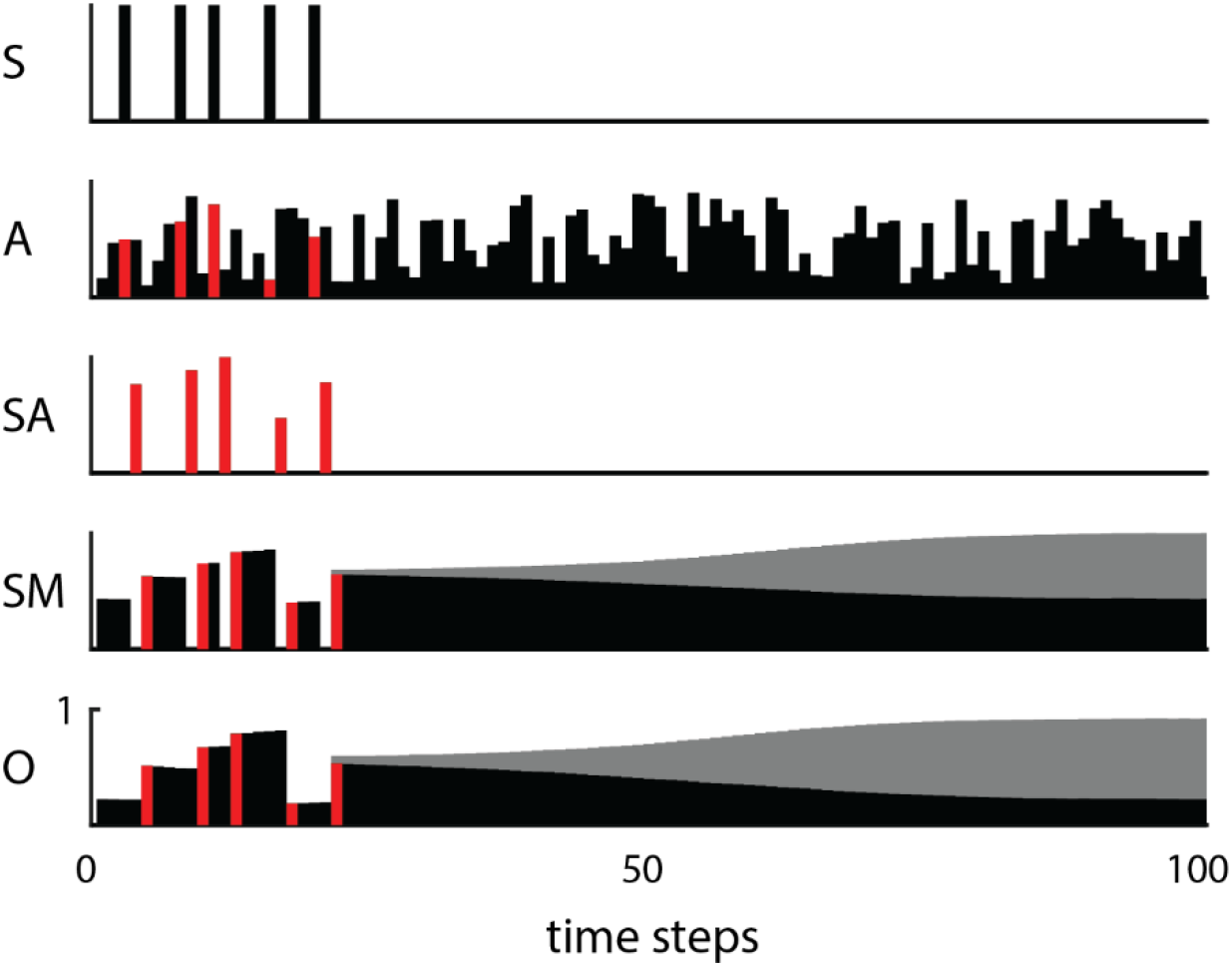
Relaxation to attractors occurs during long delays after the cue (S). Convergence to fixed point attractors occurs in both hidden units (SM) and output units (O) depending on the level of the sampled value (SA which is obtained as a non-linear combination of the cue and the analog value (A) at the time of the cue).

We investigated the attractors for reduced networks with five structurally different SAH network architectures with different connectivity constraints. Figure 10 plots the evolution of the value of a storage hidden unit in the time-steps after the last sample as a function of the last stored value. As time-steps increase, the stored values asymptote to one of two attractor states, depending on initial value.

**Figure 10.**
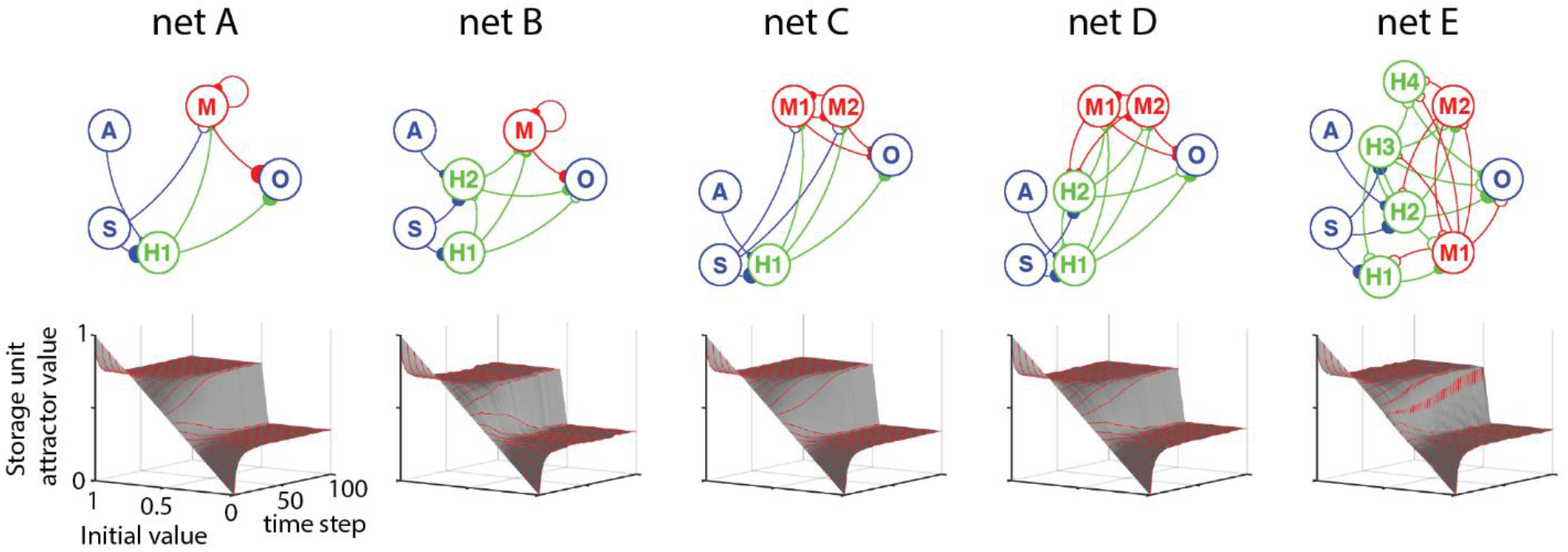
Different network architectures with the same unit activation functions exhibit similar relaxation dynamics. Input and output units are drawn in blue, gating units (H) in green, and storage units (M) in red. Excitatory connections drawn with solid boutons, inhibitory units with open boutons. Bouton size represents connection strength. 3D plots show activity values of a storage hidden unit as a function of initial value and time-steps without renewed input.

Surprisingly, despite the significant differences in network architecture, the five SAH networks exhibited very similar relaxation characteristics in terms of attractor value, number of attractors and time course of relaxation. These findings suggest that relaxation to fixed-point attractors in SAH networks is relatively independent of network architecture.

We found that the SAH network fixed-point attractor depended on the sigmoidal input-output function of the units. Figure 11A plots the value of a storage hidden unit as a function of time and initial value for 3 different sigmoid curves. As shown, the slope of the sigmoid function (“temperature” T) determined whether there was one attractor, independent of the last stored value (T=0.75), or two attractors, depending on the last stored value. The slope of the sigmoid activation function altered the number and values of fixed points similarly for all the five SAH network architectures (Figure 11B).

**Figure 11.**
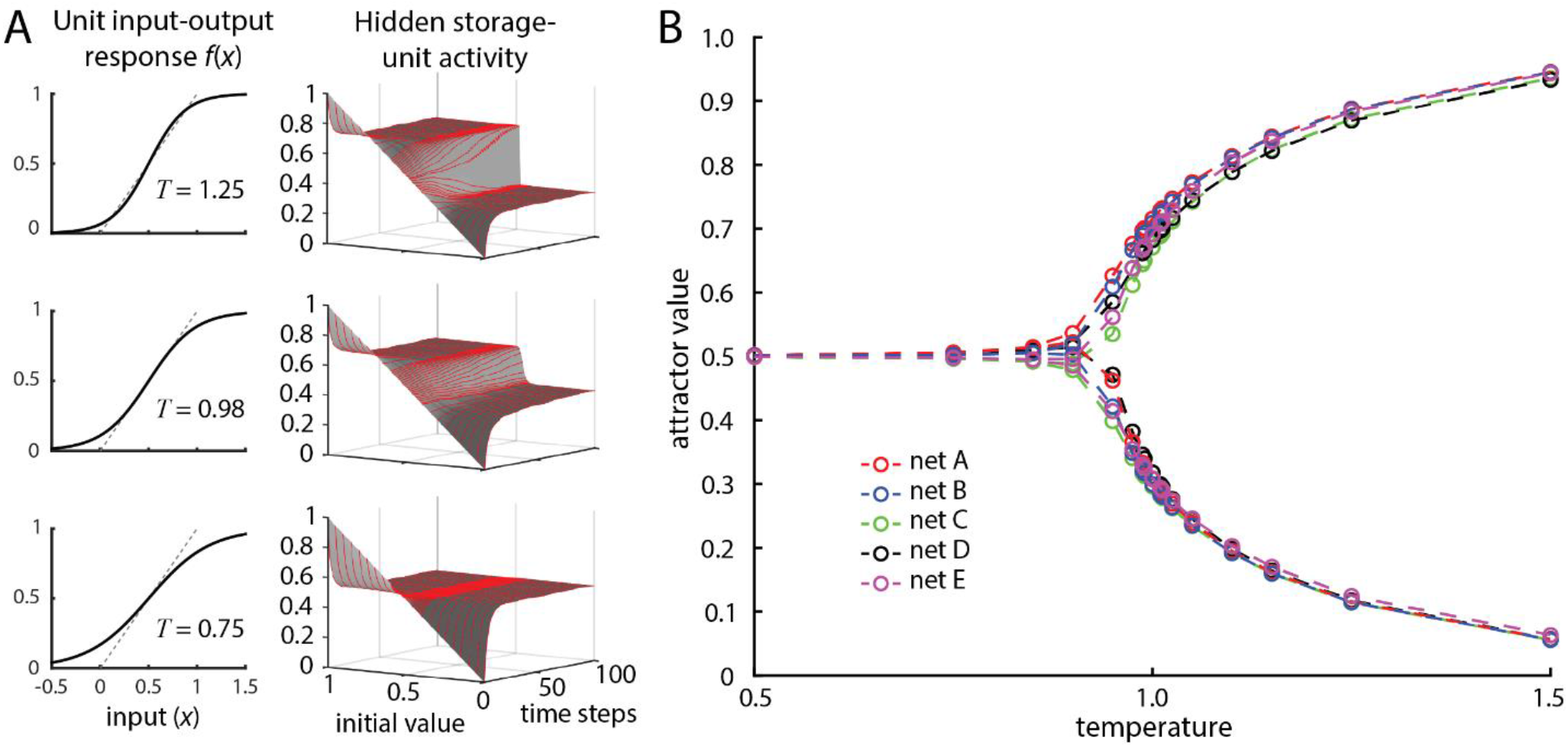
A) Fixed-point attractors vary in number and value with the slope (T for “temperature”) of the unit input-output (activation) function. B) Mean terminal attractor values of hidden storage-units follow a similar bifurcation pattern for the different network structures shown in figure 10.

The attractor values for dual input networks were explored in a similar manner: special input patterns were created which held Gate, Signal1, and Signal2 inputs active for a single time-step and then held all future inputs at zero for the next 100 time-steps. For a given network the signal input-levels were varied independently from 0.0 to 1.0 in steps of 0.1 to yield a 11×11 grid of input test patterns. The network output values for each of these test patterns were taken at the 100 time-step mark as the network’s attractor values. These simulations used temperature T= 1 for all units. Figure 12 shows how these attractor values change over the grid of input patterns for each output at common mode levels of 0%, 30%, 50%, and 60%. Most attractor values were low for a correspondingly low sampled value or were high at a high sampled value, with a transition point at a middle sample value, as observed in attractor behavior of single input SAH networks. At a common mode of 0%, the Output1 and Output2 attractor values were largely independent except at the transition point which could be influenced somewhat by the other input. At 30% the attractor value for Output1 was very slightly modified by that for Output2. This is seen as a small step in the attractor level for low Sample1 values as the Sample2 value passes the transition point for Output2. At 50% this step becomes larger, and the attractor values are more dependent on both samples. This dependence increases as the transition line becomes more diagonal. At 60% this dependence is much more pronounced as the correlation between the Ouput1 and Output2 attractor values becomes very high, and the transition point correlates well with the mean of the two sample values. The differences between these attractor maps seem to provide information on how much overlap there is between the subnetworks for each output, and hence give an estimate on the common mode value for which they were trained.

**Figure 12.**
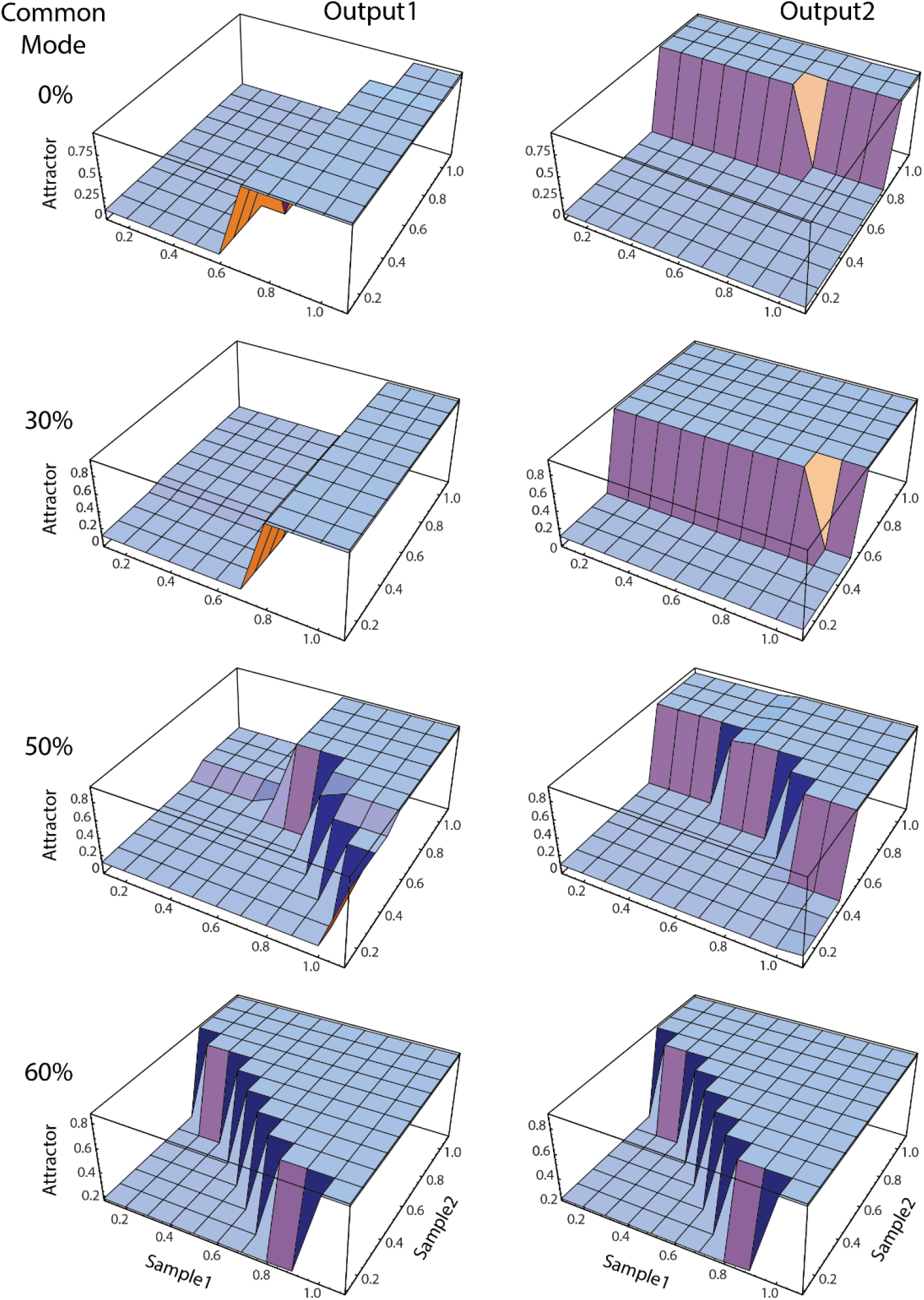
Attractor values when holding the sampled values for Signal1 and Signal2 for 100 time-steps on networks trained with various levels of common mode before any weight or unit reduction. The attractors for Output1 and Output2 become more interdependent as the common mode of the training set increases.

## Discussion

These simulations have elucidated the mechanism of memory storage in recurrent neural networks that implement a SAH function. When two inputs with common mode signal are converted to two corresponding remembered outputs, the networks trained on signals with appreciable common mode typically exhibit hidden units that process both signals. Such dual-input networks are relevant to understanding biological mechanisms mediating storage of multiple sensory signals. For example, sensory inputs from the two ears or two eyes typically have components in common, and many cortical units respond to inputs from either side (Kandel 2021). The differences in the separate signals underlie sound localization and stereopsis. Similarly, the inputs from Ia spindle afferents of a muscle have considerable common mode (Blum 2020; Matthews 1981) and converge on common postsynaptic cells in spinal cord and brainstem. The degree to which the brain exploits the large amount of common mode for memory storage remains to be investigated. It seems reasonable to assume that memory of sensory events would include memory for the activity of the neurons that carry common mode signals. However, the difference between two inputs can also be remembered, as in memory for sound localization or stereopsis. This would seem more easily performed if the difference is explicitly coded in higher-order neurons.

Our networks were not required to calculate the common mode signals in the two input signals (for 0 < alpha < 1), so the only exact solutions these networks can produce must involve two separate pathways, one for each signal/output pair. However, separate subnetworks only seem to occur for low to moderately low amounts of common mode. Networks trained with intermediate common mode values most frequently have some overlap between both input samples and one, or both, of the outputs. The point at which this transition occurs depends on several factors including the length of the training epoch and the number of training iterations. For example, at 50% common mode it can take more than 50,000 training iterations with epochs of 100 steps for a network to learn to create separate solutions for each signal/output pair. However, for many networks with significant common mode and moderate amounts of training, the networks take an average of the input samples and then subtract out contributions of opposing samples to generate the correct separate output values. There are several reasons why this could happen. First, for significant common mode the sample average is a good first-order estimate of the output values and becomes a better estimate with increased proportions of common mode. This mechanism would also scale well for networks dealing with a larger number of signals. Second, once the network has learned this approach, it may become stuck in a local minimum for the gradient-descent training algorithm. This becomes more likely as the network is reduced and there are fewer degrees of freedom to work around local flat spots in the gradient-descent error-space. Lastly, the network topology and initial connection strengths with excitatory-only and inhibitory-only hidden units may encourage the sums and differences of averaged activity to utilize all the network resources available.

In large scale classification networks, models can be retrained on adversarial inputs to improve performance against misclassification of inputs that are nearby in input space (Durán 2023). One could consider our Signal1 to be an adversarial input to the network for Output2, and the same for Signal2 versus Output1. Any non-canceling use of an input for its opposing output can only increase the error for that output on average. This is somewhat different from the typical definition for an adversarial input which would cause the network to misclassify the entire input. In addition, both inputs are presented simultaneously rather than as separate instances. However, the networks presented here with low amounts of common mode can reach lower error values in fewer training iterations than networks with no common mode.

The dynamic operation of 2D SAH networks can be effectively visualized through animations. Examples are shown on the website: https://depts.washington.edu/fetzweb/neural-networks.html. Video 2 illustrates the operation of the SAH network in Figure 1A, with activities scrolling through time. Videos 3 and 4 show the operation of a reduced 2D network with sinusoidal and pulse inputs. Video 5 shows how a lesion of a single hidden unit in a 2D network can selectively affect one of the outputs.

Our networks used units with analog activity between 0 and 1, and have sigmoidal input-output transforms, in contrast to biological neurons that carry discrete action potentials that are triggered by summing post-synaptic potentials to threshold. However, the solutions obtained for the analog units can be converted to networks of spiking integrate-and-fire units that perform the same functions by replacing each continuous unit with a set of corresponding spiking units (Maier, Shupe & Fetz, 2003). This conversion involves replacing the simple symmetric sigmoidal functions with custom sigmoidal functions that capture the input-output function of integrate-and-fire units. The errors initially produced by this substitution are then eliminated by retraining. The success of this transformation indicates that the proper interpretation of the analog activity of our units is that it represents the average rate of a population of spiking neurons.

## Future directions

Some of the variation between networks trained with different levels common mode can be attributed to the distribution of the input signal values and non-linear shape of the logistic activation function, where output values close to 0.0 and 1.0 require large inputs. Future networks could use additional randomization to normalize input signal variance, and limit output target values to the more central region of the activation function. Another area for future exploration involves networks without the restriction on self-connections and recurrent connections within the inhibitory group.

## Acknowledgements

This work was supported by NIH grants NS12542 and RR00166.

